# Adaptive Optics Rolling Slit Ophthalmoscope: Combining cellular-resolution, high-speed and large field-of-view in a multimodal retinal imager

**DOI:** 10.1101/2024.12.23.630080

**Authors:** Léa Krafft, Pierre Senee, Ana Alexandra Brad, Michael Atlan, Michel Paques, Olivier Thouvenin, Pedro Mecê, Serge Meimon

**Affiliations:** DOTA, ONERA, Université Paris Saclay F-91123 Palaiseau, France; Quantel Medical, Cournon d’Auvergne, France; Institut Langevin, ESPCI Paris, CNRS, PSL University, Paris, France; Centre d’Investigation Clinique 1423, Quinze-Vingts National Ophthalmology Hospital, DGOS, INSERM, Paris, France

## Abstract

Label-free optical imaging systems capable of monitoring dynamic biological processes over a large field-of-view (FOV) are essential for advancing our understanding of retinal and neurovascular health. However, existing imaging modalities often involve trade-offs between spatial resolution, contrast, FOV, and frame rate. In this study, we present the Adaptive Optics Rolling Slit Ophthalmoscope (AO-RSO), a novel imaging system that integrates the high-speed, wide-FOV capabilities of camera-based systems with the enhanced contrast and multimodal functionality of point-scanning techniques—without compromising resolution or speed. The AO-RSO utilizes line illumination synchronized with the rolling shutter of a high-speed sCMOS camera, enabling precise and dynamic spatial selection of detected photons: back-scattered photons for near-confocal bright-field imaging and forward-scattered photons for phase-contrast imaging of translucent retinal features. This system successfully visualizes diverse retinal structures, including cone and rod photoreceptors, nerve fiber bundles, red blood cells, vessel walls, and ganglion cells, across a wide retinal area (4.5°× 2.5°). With a large FOV and frame rates up to 200 Hz, the AO-RSO enables the quantification of blood flow, tracking of red blood cells within hundreds of capillaries, and evaluation of thousands of photoreceptors in the living human retina. This capability opens new opportunities for functional neuronal imaging, neurovascular coupling studies, and the early detection of retinal and neurodegenerative diseases.

## 1. Introduction

Recently, there has been growing interest in developing label-free optical imaging systems capable of monitoring the dynamic processes of biological tissues, such as blood flow, cellular activity, and metabolism, and their interrelationships in applications such as neurovascular coupling [1–5]. To achieve this, imaging techniques must offer sufficient spatial resolution and contrast to distinguish individual cells and biological features, as well as high temporal resolution to track dynamic processes accurately. Furthermore, large field-of-view (FOV) imaging is essential for monitoring diverse cell populations in-vivo, ideally capturing thousands of individual biological structures simultaneously. Meeting all these requirements presents a significant technical challenge [1, 6, 7]. Although camera-based systems excel in capturing dynamic phenomena over large areas at high speeds, they lack the inherent ability to reject multiply scattered photons, which limits image contrast. Conversely, laser-scanning modalities offer better photon rejection and higher contrast but suffer from low pixel throughput, restricting their ability to visualize cellular dynamics across wide FOVs.

This challenge is particularly critical in in-vivo human retinal imaging, a translucent organ surrounded by thick scattering tissues, because of the constant eye motion moves and presence of dynamic optical aberrations [8–10]. The retina, due to its unique optical properties, is the only part of the central nervous system that can be visualized non-invasively in-vivo, making it an invaluable target for studying neuronal and vascular activity directly in humans and related retinal and neurodegenerative diseases [4, 11]. By incorporating adaptive optics (AO), it is possible to correct ocular aberrations in real time, enabling cellular resolution imaging of the retina [10, 12, 13].

Historically, the first AO-based ophthalmoscope was a fundus camera, using a 2D detector, known as the AO flood illumination ophthalmoscope (AO-FIO) [12]. However, due to the low image contrast and limitations of early camera technology, researchers shifted to point-scanning techniques, such as the AO Scanning Laser Ophthalmoscope (AO-SLO) [14]. Initially, AO-SLO was employed to capture back-scattered photons for imaging highly reflective retinal features, such as photoreceptors and nerve fibers in a confocal bright-field mode. Later, the introduction of lateral displacement between the illumination and detection paths allowed the rejection of back-scattered photons and the detection of multiply-scattered photons forward-scattered by translucent retinal structures, producing phase-contrast images [15–18]. This mode enabled the visualization of translucent features, such as red blood cells, vessel walls, ganglion cells, photoreceptor inner segments, and immune cells [15, 17–22]. By integrating confocal and phase contrast imaging, AO-SLO has become a powerful multimodal imaging tool, providing complementary details that have significantly improved our understanding of retinal structure and function. Despite these advances, AO-SLO is inherently constrained by its low pixel throughput. Quantifying dynamic processes, such as blood flow, often requires sacrificing FOV to achieve the necessary temporal resolution [23]. Furthermore, the slow speed of mechanical scanners introduces motion artifacts from involuntary eye movements, distorting retinal images, and compromising the precision of quantitative assessment derived from the data [24]. Recent advances in camera technology have led to the exploration of high-contrast retinal imaging systems capable of achieving wider FOVs or high frame rates.

An early attempt to increase imaging speed was the AO Line-Scanning Ophthalmoscope (AO-LSO) [25]. Using this technology, Gu et al. achieved frame rates of 200 Hz within a narrow 1.2°× 1.2° FOV, allowing the quantification of blood flow in a few capillaries [26]. However, the limited FOV prevented long-term monitoring of blood flow, as the capillaries would often drift out of view due to eye movements during acquisition. Furthermore, AO-LSO methods have mainly focused on near-confocal imaging of back-scattered photons, without incorporating phase contrast, leading to noisy blood flow measurements [27].

Recently, other groups combined AO-FIO with patterned illumination based on digital micromirror devices (DMD) [27–32]. Initially, DMDs were used to enhance bright field image contrast and resolution on wide FOVs, albeit at the cost of imaging speed [29, 30]. Later, DMD was used to project complementary illumination patterns, enabling multimodal imaging with bright-field and phase contrast modes through post-processing by selecting illuminated and non-illuminated areas respectively [27, 28, 31, 33]. However, this approach requires multiple acquisitions and suffers from significant light loss due to the DMD, necessitating image averaging to improve signal-to-noise ratio (SNR). These limitations result in trade-offs between imaging contrast, speed, and FOV [27, 28, 31, 33]. Finally, recent studies by our group have demonstrated, both experimentally and theoretically [30, 34], that line illumination achieves optimal imaging contrast and signal-to-noise ratio (SNR), comparable to point-scanning illumination, for biological samples, including the retina.

Here, we present the AO Rolling Slit Ophthalmoscope (AO-RSO), a camera-based imaging system that uniquely combines the speed and distortion-free wide FOV of camera-based modalities with the enhanced contrast and multimodal capabilities of point-scanning techniques - all without trade-offs. Building on our prior findings on the benefits of line illumination, the AO-RSO employs a Powell lens to focus light into a line pattern, which is synchronously scanned with the rolling shutter of a high-speed sCMOS camera. The rolling shutter acts as a moving slit, enabling spatial selection of photons and the generation of different contrast modes. A defining feature of the AO-RSO is its real-time digital switching capability, allowing precise adjustments to line-pixel offsets and exposure time to optimize image contrast. With the AO-RSO, we demonstrate high-contrast bright-field and phase contrast imaging over a wide FOV (4.5° × 2.5° FOV) and at high frame rates (up to 200Hz), enabling key applications such as blood flow quantification, red blood cell tracking, and photoreceptor counting across a wide FOV.

## 2. Method

### 2.1 Experimental setup

The experimental setup is illustrated in Figure 1-1. We modified our AO-FIO (previously described elsewhere [13, 30, 31]) to integrate the AO-RSO imaging module. The AO-RSO imaging module benefits from correction for ocular aberration through the AO system. The AO-RSO system is composed of a superluminescent diode (SLD) that emits at 850 nm with a spectral bandwidth of 55 nm (SLD850S-A20W, Thorlabs). The SLD is collimated and shaped into a 1D line source via a Powell lens (30° fan angle, ♯43 − 473, Edmund Optics). Unlike cylindrical lenses, Powell lenses ensure uniform illumination along the line. After the Powell lens, an achromatic doublet of 30 mm focal length is introduced to form a uniform line at the focal plane F1. A series of telescopes ensure the conjugation and magnification of F1 to the retinal plane. The resulting line projected onto the retina spans a lateral field of 4.5° (Figure 1-a) and a line width of 7*μm*. The choice of line width is motivated by our previous studies, in which we demonstrated, both experimentally and theoretically, that this width is sufficient to optimize image contrast and SNR in bright-field mode [30, 34]. A 1D galvanometer mirror (6210*H*, Cambridge Technology) is conjugated to the pupil plane of the system and scans the line across the retina. A series of telescopes ensure the conjugation at the pupil plane of the system for the following elements: galvanometer, deformable mirror, the eye’s pupil, and the micro-lens array from the Shack-Hartmann wavefront sensor. The light scattered by the retina follows the reverse optical path before being reflected by a 50/50 beam splitter that separates the illumination and detection paths. The reflected light is focused by a tube lens and forms a line on the imaging camera plane. The imaging camera (ORCA-Fusion Digital CMOS Camera, Hamamatsu), with a 2304 × 2304-pixel sensor and 6.5*μ*m pixel pitch, operates in rolling shutter mode, synchronizing the electronic shutter with the galvanometer mirror for AO-RSO image acquisition. It can acquire at up to 200 fps for a 4.5° × 2.3° FOV (1800 × 960 pixel) or 100 fps for 4.5° × 4.5° FOV.

**Fig. 1.**
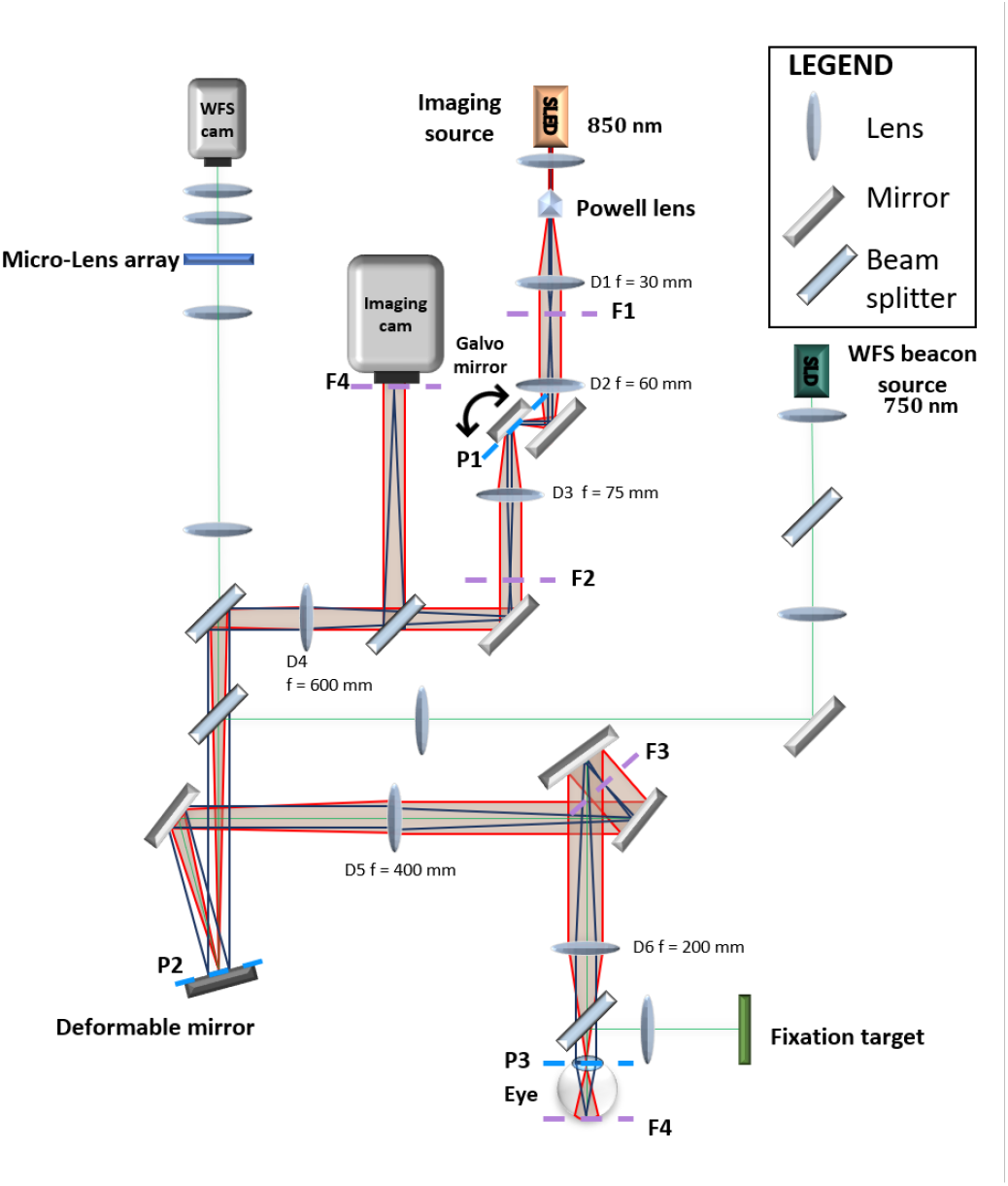
Schematic drawing of the AO-RSO system. F, P and D indicates the focal planes, pupil planes and lenses respectively. Green tracing indicates the AO light path. Red and black tracing indicates the imaging path at horizontal and vertical direction respectively.

### 2.2 Synchronization procedure for multimodal acquisition

Precise synchronization between the rolling shutter of the imaging camera and the galvanometric scanner is critical to making multimodal imaging possible. A custom Matlab code generates two synchronized digital signals composed, respectively, of a transistor transistor logic (TTL) signal to trigger the camera, along with a sawtooth signal for the galvanometer mirror. Both signals are converted to analogic signals and sent to the respective devices via a National Instruments DAQ card (PCIe-6321). The camera operates in “Edge Trigger” mode, for which a new frame acquisition begins on the rising edge of the TTL signal. This signal defines the frame rate of the imaging sequence.

As shown in Figure 2-A, the rolling shutter exposes pixel arrays sequentially, with each array exposed for a duration *t*_*array*_. The width of the slit is defined by the number of simultaneously exposed pixel arrays on the rolling shutter mode, while the interval between arrays *t*_*i*_ reflects the time the shutter needs before starting to expose the next pixel array and is set as 5*μs*. Off-axis detection is defined by the delay *t*_*offset*_, which represents the temporal offset between the galvanometric scan and the exposed pixel arrays. In other words, *t*_*offset*_ corresponds to a spatial offset of the detection aperture relative to the line illumination. All these detection parameters can be controlled via a custom-made Matlab graphic user interface (GUI). During image session, one can define the acquisition mode between bright-field and off-axis detection, with detection settings predefined as follows.

**Fig. 2.**
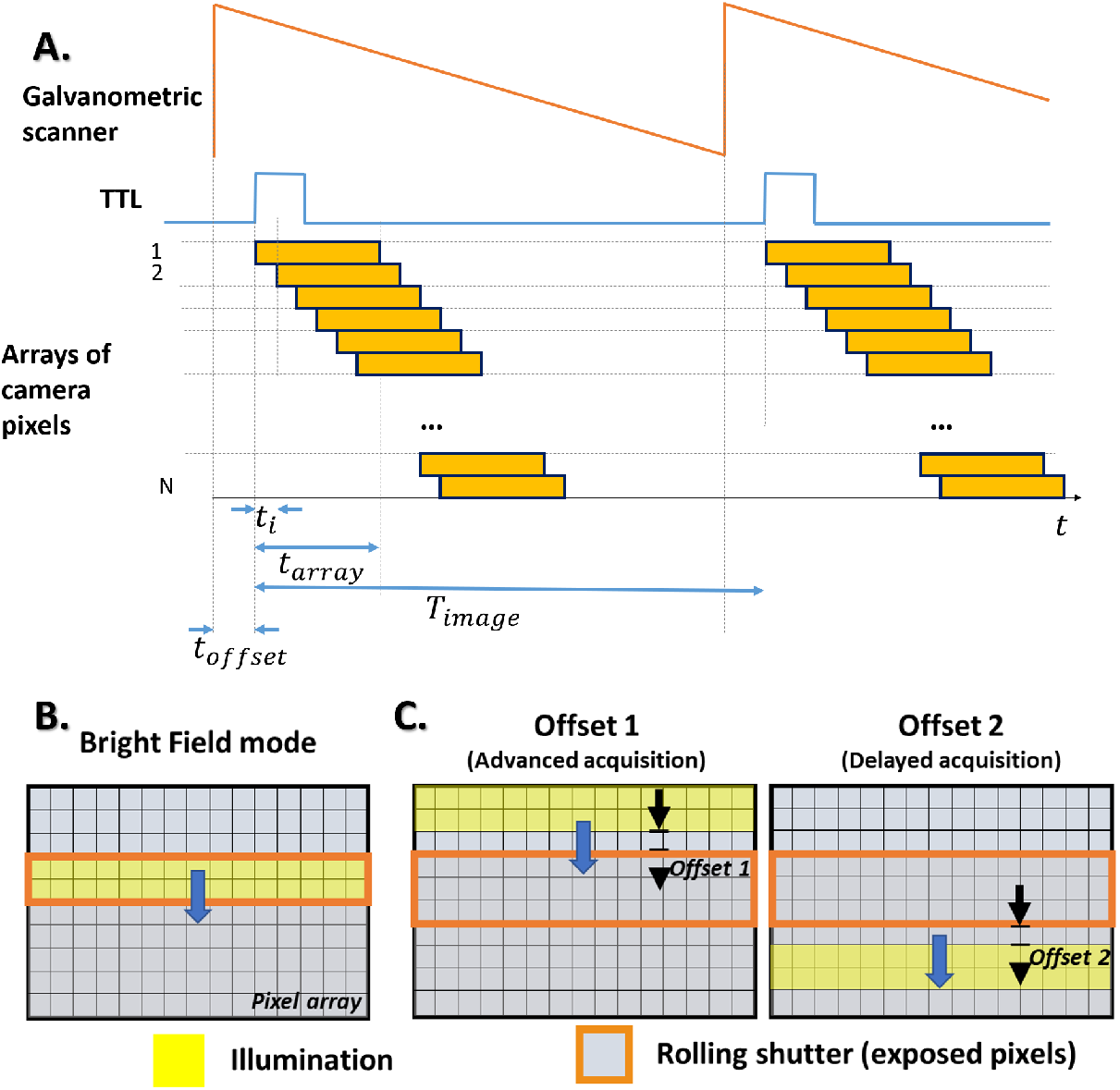
A. Chronogram of the synchronization between the galvanometer mirror and the sCMOS camera. *t*_*i*_ corresponds to the time between the exposure of two consecutive arrays of pixels. *t*_*array*_ is the exposure time of a pixel array. *T*_*image*_ is the period of the TTL signal, namely the time between the start of two consecutive frames. *t*_*offset*_ corresponds to the delay between the scan of the galvanometric mirror and the array of pixels being exposed. B) *t*_*offset*_ is equal to zero when acquiring bright-field images. C) *t*_*offset*_ is positive and negative allowing to acquire photons outside the illumination line, for offset 1 and 2 images, which combined generate phase contrast images.

For bright-field imaging mode, the line illumination and the rolling shutter are in phase, and the width of the slit is chosen to match the line width (10-pixel array, for an equivalent of 2 Airy disk diameter - ADD - aperture), resulting in *t*_*array*_ of 50*μ*s. In this configuration, most of multiply scattered and out-of-focus photons are filtered out, since only 1/100th of the field is illuminated [30], thus enabling to generate bright-field images with contrast comparable to those acquired with AO-SLO (Fig. 2-B).

By introducing an offset between the line illumination position and the exposed pixels of the rolling shutter, one can generate an off-axis detection similar to the AO-SLO. In this mode, back-scattered photons are filtered out, and multiply scattered photons responsible for phase contrast are detected. In this configuration, *t*_*offset*_ is initially set at 50*μs* (2ADD equivalent), with a slit width initially set with *t*_*array*_ = 300*μs* (12ADD equivalent). To generate phase contrast images, one needs to subtract two offset images with opposite *t*_*offset*_, meaning a line illumination in advance and delayed compared to the rolling shutter (detection of pixels on the top or on the bottom of the line position, see Fig. 2-C).

For phase-contrast images, detection parameters such as *t*_*offset*_ and *t*_*array*_ can be further adjusted during image session to optimize the image contrast for a retinal feature of interest. With the custom-made Matlab GUI, we can also adjust the detection parameters at the speed of the camera (for each frame), enabling to switch from bright-field to phase-contrast images modes sequentially in one image sequence.

### 2.3 Image acquisition

The data presented in this study were acquired in four subjects of health. All participants followed institutional guidelines and adhered to the tenets of the Declaration of Helsinki. Informed consent was obtained after explaining the purpose, procedures, and potential outcomes of the study. All imaging sequences were collected with the room lights off. The subject’s eye was cyclopleged and dilated using tropicamide 0.5%. The eye and head were aligned with the imaging system using a chinrest mounted on a motorized XYZ translation stage [35]. A fixation target composed of a yellow crosshair was used to guide the subject’s gaze and minimize eye motion during the imaging session. The AO beacon source operated at a power of 3.7*μW*, and the imaging source at 1*mW*. The illumination irradiance remained well within the maximum permissible radiant exposure specified by the ISO standards (ISO 15004-2 (2007)) for group 2 devices. For real-time monitoring and acquisition of AO-RSO retinal images, we used Holovibes software [36] combined with a Nvidia GTX Titan Xp GPU (CUDA 9) and an Active Silicon FireBird Frame Grabber (BASEBOARD 2xCXP662), which ensures smooth image rendering and high-throughput recording of raw images directly to disk without frame drops.

### 2.4 Image processing

#### Image alignment

To compensate for fixational eye movements, the acquired image sequences were aligned using custom Matlab software. The image alignment algorithm is composed of three steps. 1) rigid alignment using a custom made high spatial frequency-based phase correlation algorithm which is detailed elsewhere [37, 38]; 2) rotation correction using a masked phase correlation approach [39], and 3) a strip-based alignment with subpixel precision [40]. After image registration, we averaged around 10 images to generate bright-field images and 100 images to generate phase-contrast and motion-contrast (perfusion maps) images. Image montages were performed using ImageJ plug-in MosaicJ [41].

#### Photoreceptor density

To generate the pointwise density map, we divided the cone mosaic image into an overlapping grid of 200 × 200 pixels (corresponding to 150 *μm* × 150 *μm*) regions of interest (ROI), where each ROI was displaced from the previous by 30 pixels (corresponding to 20*μm*). These values were chosen empirically to provide a good trade-off between point-wise accuracy and map smoothness. Then, the cone density and spacing were computed for each ROI using a fully automated algorithm based on modal spacing as described in [42]. Bicubic interpolation was used to increase the size of the cone density map to match the cone mosaic image [43].

#### Perfusion Maps and blood flow assessment

Contrast motion images were generated using phase-contrast images, in two steps. First, to favor the structure of red blood cells and filter out other retinal structures and noise, we subtracted each image by its blurred version using a Gaussian filter (sigma = 5) [33]. Finally, to eliminate the contribution of static or slow retinal features, we applied a highpass temporal filter by subtracting consecutive images [33]. After these two steps, the temporal standard deviation of each pixel was computed, allowing the generation of perfusion maps. Frequency color-coded perfusion maps were generated with four different perfusion map images that underwent different temporal filtering using Matlab function bandpass. Temporal filters had the following cutoff frequencies: 1) 5-15Hz, 15-25Hz, 25-35Hz, and 35-50Hz. Each of these temporal-filtered perfusion maps had a color assigned, blue, green, yellow, and red, respectively, using the HSV space. Finally, the calculation of blood flow velocity was done by kymographs as detailed in [26, 27].

## 3. Results

### 3.1 High-Contrast Rod and Cone Imaging Across a Wide Field at High Frame Rates

The AO-RSO imaging method combines the advantages of both AO-SLO and AO-FIO modalities. When operating the AO-RSO in Bright-Field mode — where the line illumination and the camera’s rolling shutter are synchronized and in phase — most unwanted multiply-scattered photons are filtered out. This enables the acquisition of high-contrast images of the in-vivo human retina. Foveal cone photoreceptors are resolved with high contrast throughout the FOV of 4.5° × 2.5° (Fig. 3A), surpassing the typical FOV of AO-SLO systems, which is approximately 1° × 1° for an equivalent spatial sampling.

**Fig. 3.**
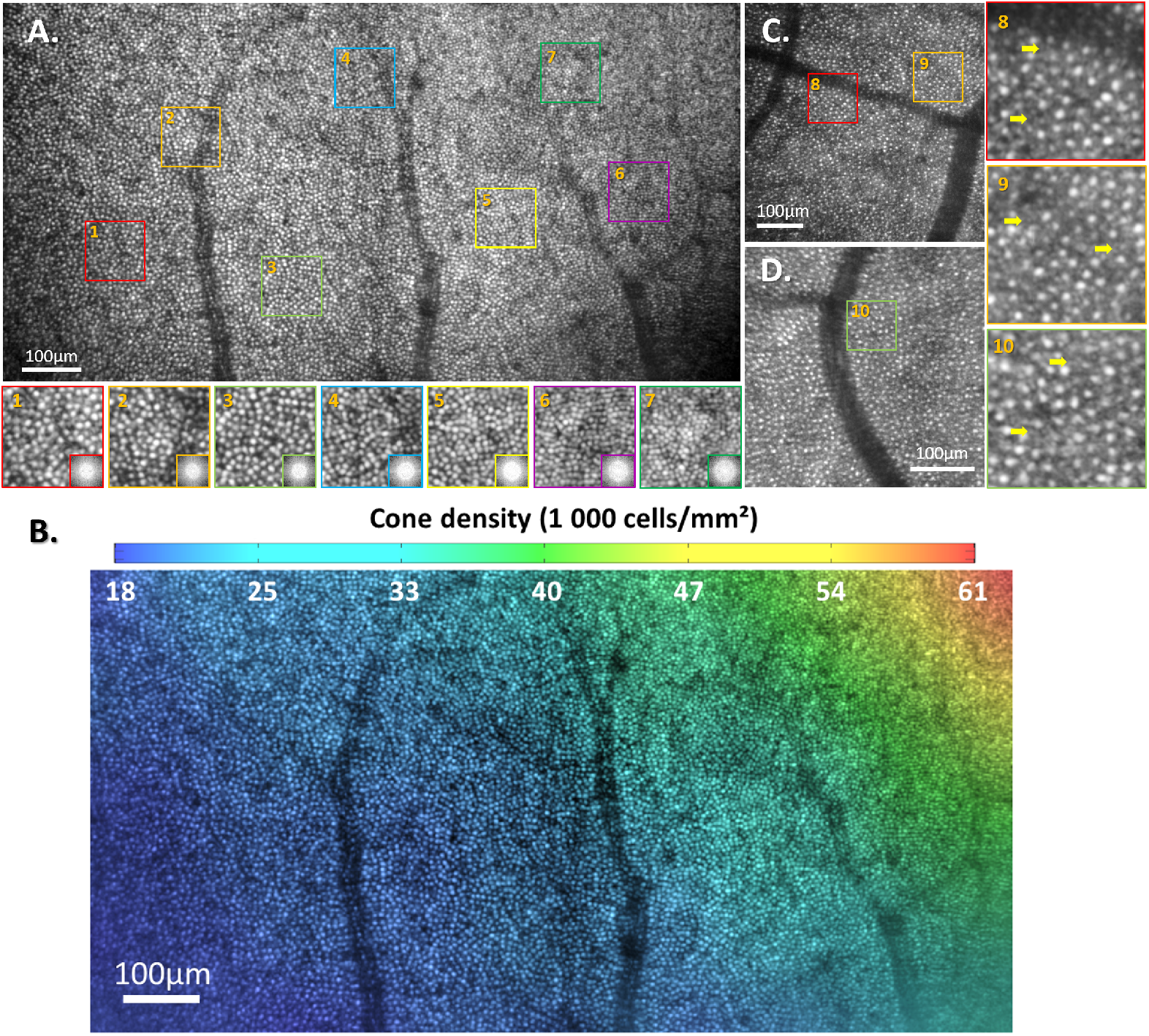
A. Image of cone photoreceptors near the fovea, highlighting the high-contrast and resolution achieved with the AO-RSO system. Zoom-in images across the FOV and their respective PSDs on the bottom right, where the Yellot’s ring can be visualized. B. Cone density distribution color coded for a 25 images averaged. C. and D. Images of photoreceptors at peripheral retina for two different subjects, where rods can be visualized. Yellow arrows indicate examples of rods in Zoom-in images).

Zoomed-in images from different regions demonstrate that high resolution and contrast are maintained uniformly across the entire FOV. This consistency is further supported by the analysis of their respective power spectral density (PSD), shown at the bottom right of the zoomed images. In particular, all PSDs exhibit the Yellott’s ring, the spectral signature of the photoreceptor mosaic [44]. Using the PSD, we can extract local cone density measurements [42], represented in a color-coded map in Fig. 3B.

The large FOV enables cone density measurements ranging from 18,000 cones/mm^2^ to 61,000 cones/mm^2^, consistent with histological data and AO-assisted cone density measurements for a healthy young subject at this eccentricity [42, 45]. Our method was able to detect thousands of photoreceptors in less than 50 ms of image acquisition duration, where a system with a typical 1° × 1° FOV would need to acquire at least 15 images at different positions (acquired at 30Hz) to obtain the same amount of information, thus greatly benefiting clinical workflow.

In addition to acquiring high-resolution, high-contrast images over a large FOV, the AO-RSO method achieves a high frame rate (200 Hz in this study), enabling the generation of images free from motion-induced distortions. **Visualization 1** presents the photoreceptor image sequence from Fig. 3A. The absence of image distortion over time, along with consistently high contrast and signal-to-noise ratio (SNR) in individual frames, can be noticed as single photoreceptors can be tracked in time across the entire FOV without jitter.

The high contrast and SNR observed in single frames are achieved even though the line illumination width is approximately 2 Airy Disk Diameters (ADD). This aligns with the findings of Krafft et al. [30], who reported that a illumination pattern that cover around 1/100 of the FOV (equivalent of a line width below 2 ADD) is sufficient to achieve image contrast and SNR comparable to those of AO-SLO. Interestingly, despite the 2 ADD line width, optical sectioning does not appear to be compromised or less effective than in AO-SLO (vessels above are not visible, only their shadows). In Fig. 3A, photoreceptor cells remain visible even in areas shadowed by out-of-focus superficial blood vessels.

The ability of AO-RSO to capture fine reflective structures in the retina is further demonstrated at peripheral eccentricities, where rods surrounding cone photoreceptors are successfully imaged in two different subjects (Figs. 3C and D).

### 3.2 High-Throughput Multimodal Retinal Imaging

By actively and automatically adjusting the position of the line illumination relative to the rolling shutter, the AO-RSO system enables multimodal imaging (see Methods). Switching between Bright-Field, Offset 1, and Offset 2 imaging modes allows the acquisition of complementary retinal views with high throughput and large FOV. Figure 4 highlights the multimodal capabilities of AO-RSO in imaging the nerve fiber layer (NFL) in three subjects at different eccentricities.

**Fig. 4.**
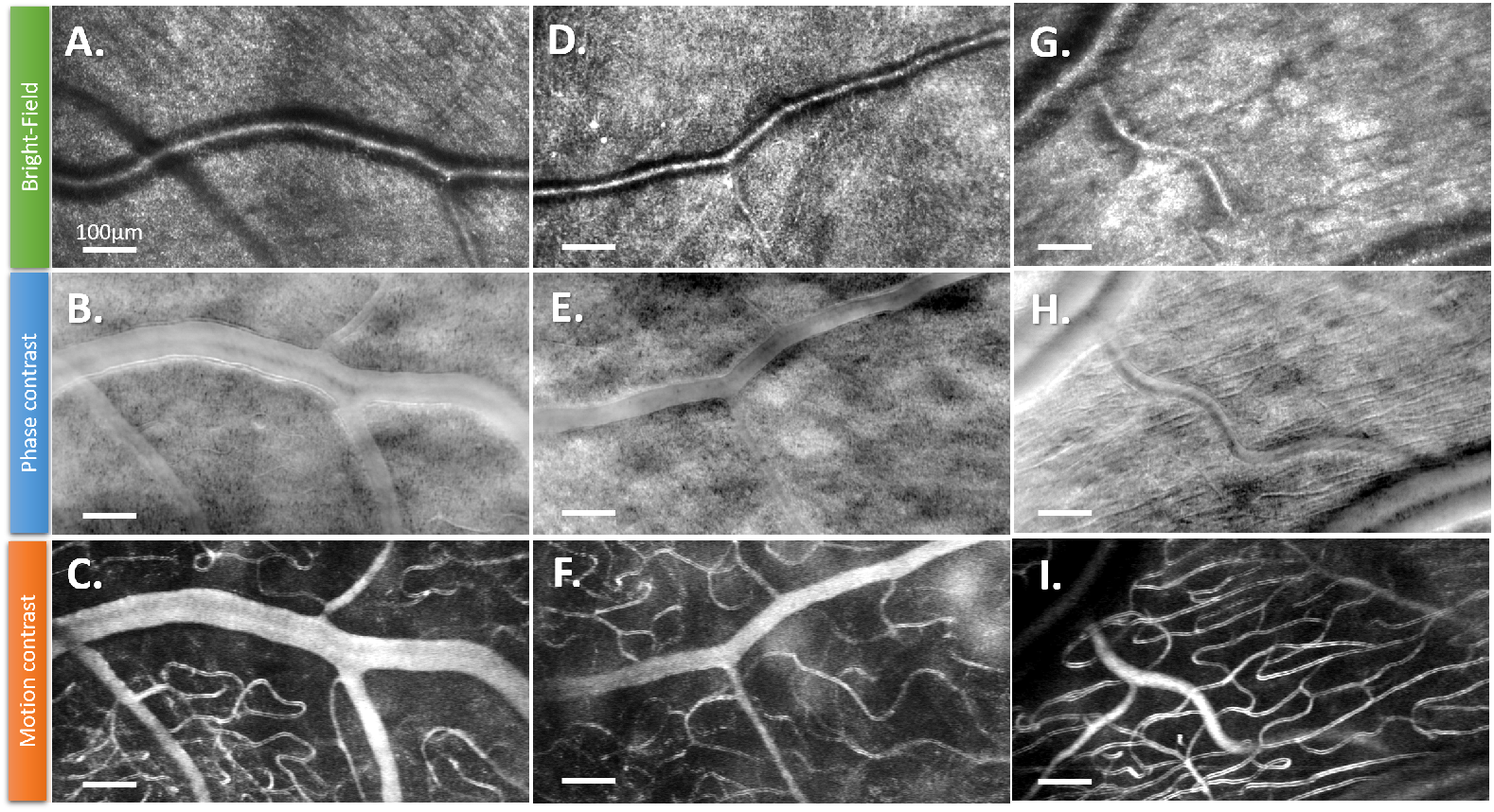
Demonstration of multimodal imaging on three different subjects (S1: A,B,C; S2: D,E,F; S3: G,H,I) when imaging at the level of the nerve fiber layer. A., D. and G. are bright-field images. B., E. and H. are phase-contrast images. C., F. and I. are motion contrast or perfusion map images.

When the line illumination and the camera rolling shutter are in phase, ballistic photons back-scattered by reflective retinal features are detected, allowing visualization of nerve fiber bundles (Fig. 4 A, D, and G). By introducing an offset between the line position and the rolling shutter, and subtracting two opposite offsets, phase-contrast images are generated (see Methods), similar to those produced by off-axis detection in AO-SLO [15, 18, 21]. Phase contrast facilitates the visualization of translucent retinal features by rejecting back-scattered photons and detecting multiply-scattered photons that back-illuminate by these structures [15, 16]. In this mode, nerve fibers disappear, while capillaries, vessel walls, and ganglion cells become visible, similar to off-axis detection in AO-SLO but with a wider FOV and higher frame rate (Fig. 4 B, E, and H).

Using a time series of phase contrast images, the standard deviation of each pixel can be calculated over time to generate motion contrast images or perfusion maps [15]. These motion-contrast images reveal small capillaries and larger vessels with high resolution and contrast, providing a detailed view of the retinal vascular network (Fig. 4 C, F, and I).

Unlike other camera-based multimodal imaging techniques that rely on DMD to project illumination patterns [27, 31], the AO-RSO method eliminates the need for extensive digital post-processing and enables to optimize the photon budget. These features are particularly advantageous in clinical settings, as it enables real-time visualization of the retina in the desired imaging mode during eye examinations.

### 3.3 Tailoring Off-Axis Detection to Optimize Phase Contrast Imaging

A key advantage of the AO-RSO method in phase-contrast imaging is the ability to adjust the off-axis detection design during an imaging session. Specifically, one can reconfigure 1) the offset between the line illumination and the slit detection (by delaying the line scan relative to the rolling shutter) and 2) the width of the slit detection (by adjusting the exposure time of a pixel array). This flexibility allows for tailoring the off-axis detection scheme to optimize phase contrast for a given translucent retinal feature. Different retinal features require distinct off-axis detection configurations due to variations in refractive index, size, shape, and depth within the retina [17, 18].

Figure 5 illustrates the importance of tailoring the off-axis detection scheme to optimize phase contrast to specifically detect a retinal feature. Fig. 5-A shows a parametric study on photoreceptor inner segment imaging, where the Michelson contrast is measured in images acquired from the same region with different slit widths, while maintaining an offset of 2 ADD. The contrast of the photoreceptor inner segment increases with slit width, peaking at 16 ADD, after which it begins to decrease. Due to the optimized image contrast, other features become visible, such as rods (green arrows) and subcellular features localized within the photoreceptor inner segment (yellow arrows), which were also observed in a previous study using a fixed designed off-axis detection AO-SLO [46].

**Fig. 5.**
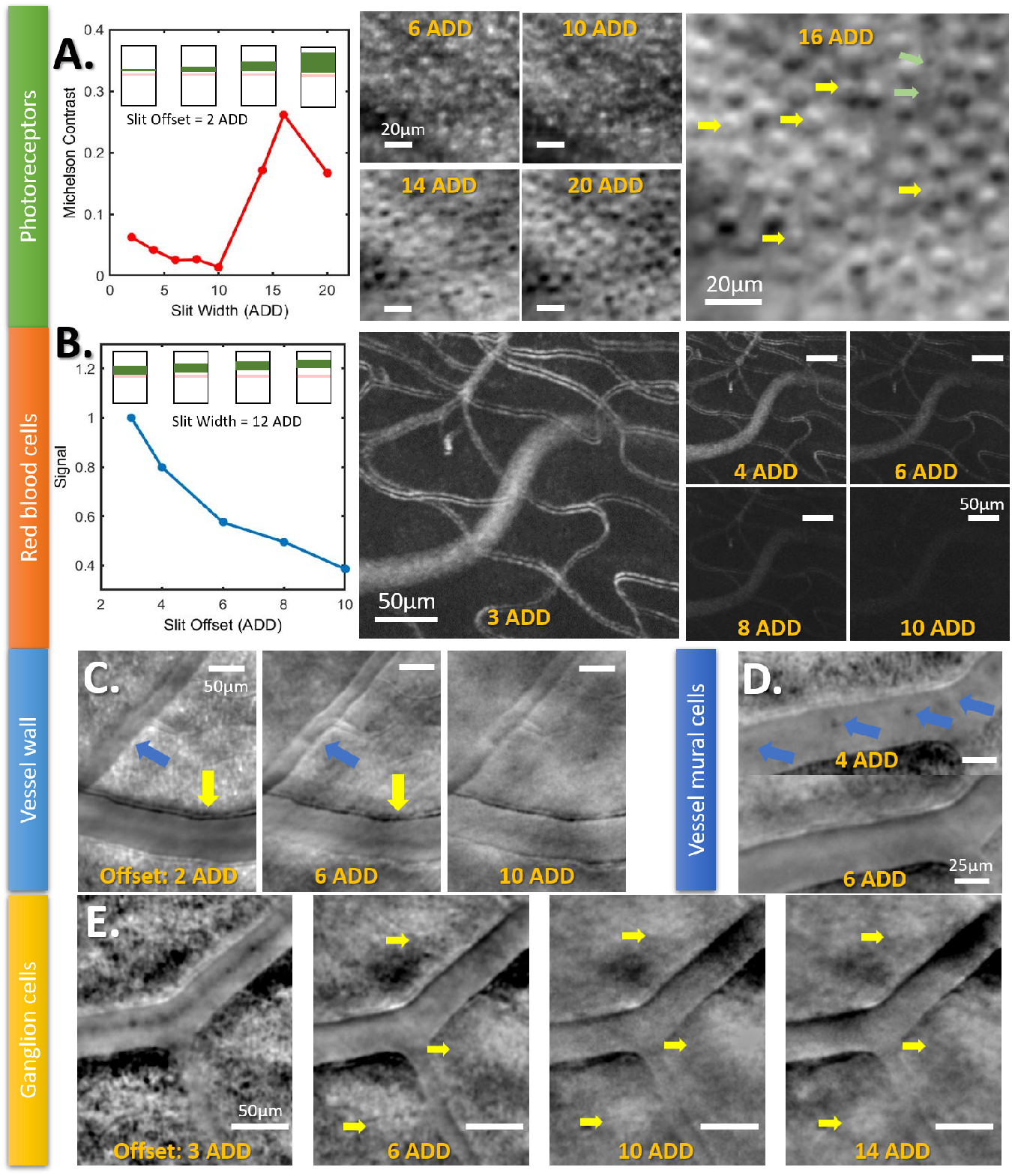
Capability to tailor off-axis detection to enhance phase contrast imaging for a retinal feature of interest. A. Phase-contrast cropped images of photoreceptor inner segment for a slit offset of 2ADD and different slit widths. The maximum contrast is reached for a slit width of 16ADD. Green arrows and yellow arrows indicates rods and subcellular cone photoreceptor features respectively. B. Motion contrast cropped images for a slit width of 12 ADD and different slit offsets. A maximum signal of motion contrast is reached for slit offset of 2ADD. C. Vessel walls at slight different depths (indicated by yellow and blue arrows) get sharper for different slit offsets, one for 2ADD and the other for 6ADD. D. Putative vessel mural cells become visible (blue arrows) for a short slit offset range, as they appear for 4ADD and disappear for 6ADD. E. Cropped images acquired at the ganglion cell layer for different slit offsets, where individual ganglion cells become visible (yellow arrows) for larger slit offsets, ranging from 10ADD to 14ADD.

A similar parametric study was conducted to improve the contrast of red blood cells (Fig. 5-B). In this case, image sequences were acquired in the same NFL region for different slit offsets, while maintaining a slit width of 12 ADD. Instead of directly evaluating red blood cell contrast, we assessed the overall motion contrast signal, as higher red blood cell contrast corresponds to a higher standard deviation over time, thus improving the SNR for motion contrast perfusion maps. As shown, the optimal contrast for a 12 ADD slit width is achieved with a minimal offset of 2 ADD. As the offset increases, the perfusion map signal dramatically decreases.

For other retinal features, different off-axis slit configurations may be required to optimize phase contrast or simply visualize the feature of interest. For example, in the case of vessel walls, Fig. 5-C demonstrates that the optimal offset for two arteries, located at slightly different depths, varies: 2 ADD for the artery indicated by the yellow arrow and 6 ADD for the artery indicated by the blue arrow. In some cases, specific cells are only visible in certain off-axis configurations, such as putative vessel mural cells in Fig. 5-D. These cells (blue arrows) are visible with a 4 ADD offset (12 ADD slit width), but disappear with a 6 ADD offset configuration.

While most retinal features in this study exhibit optimal contrast at lower offset values, retinal ganglion cells (Fig. 5-E) require larger offset values for optimal contrast. Ganglion cells, ranging from 6*μm* to 14*μm* in size, become visible with offset values between 10 and 14 ADD (yellow arrows). This finding is consistent with previous studies using multi-offset AO-SLO [21].

### 3.4 High-Throughput Imaging of Retinal Vasculature

Due to its high throughput, the AO-RSO imaging method enables, for the first time, high-resolution phase-contrast imaging over a wide FOV and at a high frame rate. This unique performance can greatly facilitate the clinical diagnosis of retinal disorders in the early stages, where high frame rate and large FOV images are necessary to capture global and local changes, and to improve patient comfort and session flow.

Figure 6 demonstrates two ways in which large FOV phase-contrast images can be explored. Figure 6-A shows a composite perfusion map, generated using motion contrast, where the color indicates images acquired at different retinal depths. A total of 9 sequences of 250 phase contrast images, acquired at 100 Hz with an axial step of 15*μm*, were captured, totaling less than 30 seconds. As previously shown, the depth of focus in phase contrast imaging of a dilated eye is sufficient to distinguish all three capillary plexuses [20, 47]. Here, even near the fovea, where the spacing of the plexus is minimal [48], the 3D organization of the capillaries and the interconnecting capillaries between the plexuses (yellow arrows) are clearly visible.

**Fig. 6.**
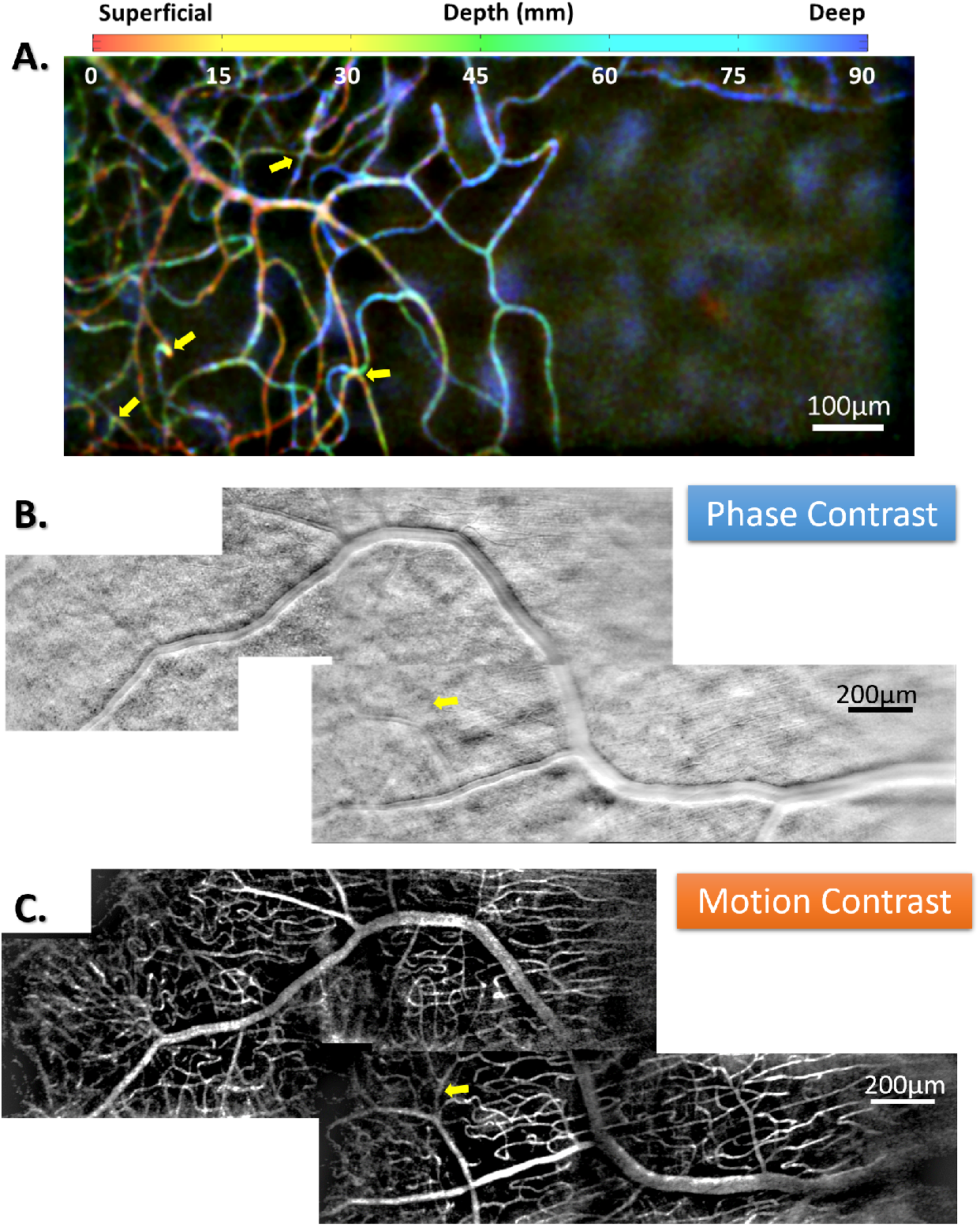
A. Composite image of 9 different perfusion maps acquired at different retinal depths (15*μm* spacing), color-coded, from red to blue for superficial to deeper plexuses at the fovea center. Arrows highlights interconnecting capillaries between plexuses. B. and C. Image montage composed of 5 images using phase-contrast and motion-contrast respectively. Yellow arrow indicates a vertical vessel not visible in phase contrast but visible with motion contrast.

Another way to leverage the high throughput of the AO-RSO system is to further expand the FOV by capturing images at multiple eccentricities and creating image montages. Figures 6-B and C show montages of 10°× 4° FOV using phase-contrast and motion-contrast images, respectively. Only five sequences of 250 images (acquired at 100 Hz) were required, resulting in a total acquisition time of less than 15 s. With high contrast of vessel walls and perfused vessels, these montages can be used to evaluate important retinal biomarkers, such as the lumen-to-wall ratio of the vessel and the perfused areas, which are critical for the evaluation of retinal diseases [49–51].

### 3.5 Blood flow visualization and quantification over a large FOV

Another important application that takes advantage of the high-throughput capability of AO-RSO in phase contrast imaging is the direct visualization and quantification of red blood cells (RBCs) within the retinal microvasculature. This feature enables, for the first time, the assessment of blood flow across a large field of view. The ability to quantify blood flow over an extended FOV offers a unique advantage for the in-depth examination of vascular diseases over time in humans [26]. Figure 7 provides an example of RBC visualization and blood flow quantification. Figure 7-A illustrates the ability of AO-RSO to directly visualize and track individual RBCs over time (indicated by arrows). RBCs exhibit a biphasic contrast, characteristic of phase contrast imaging, similar to that observed in photoreceptor imaging (Fig. 5-A). By plotting a time profile perpendicular to a capillary, the RBCs flowing in the capillary can be tracked (Fig. 7-B, top). This enables calculation of the RBC frequency (or rate or cells per second), which is color-coded in the figure. The frequency ranges from approximately 7 Hz to 37 Hz.

**Fig. 7.**
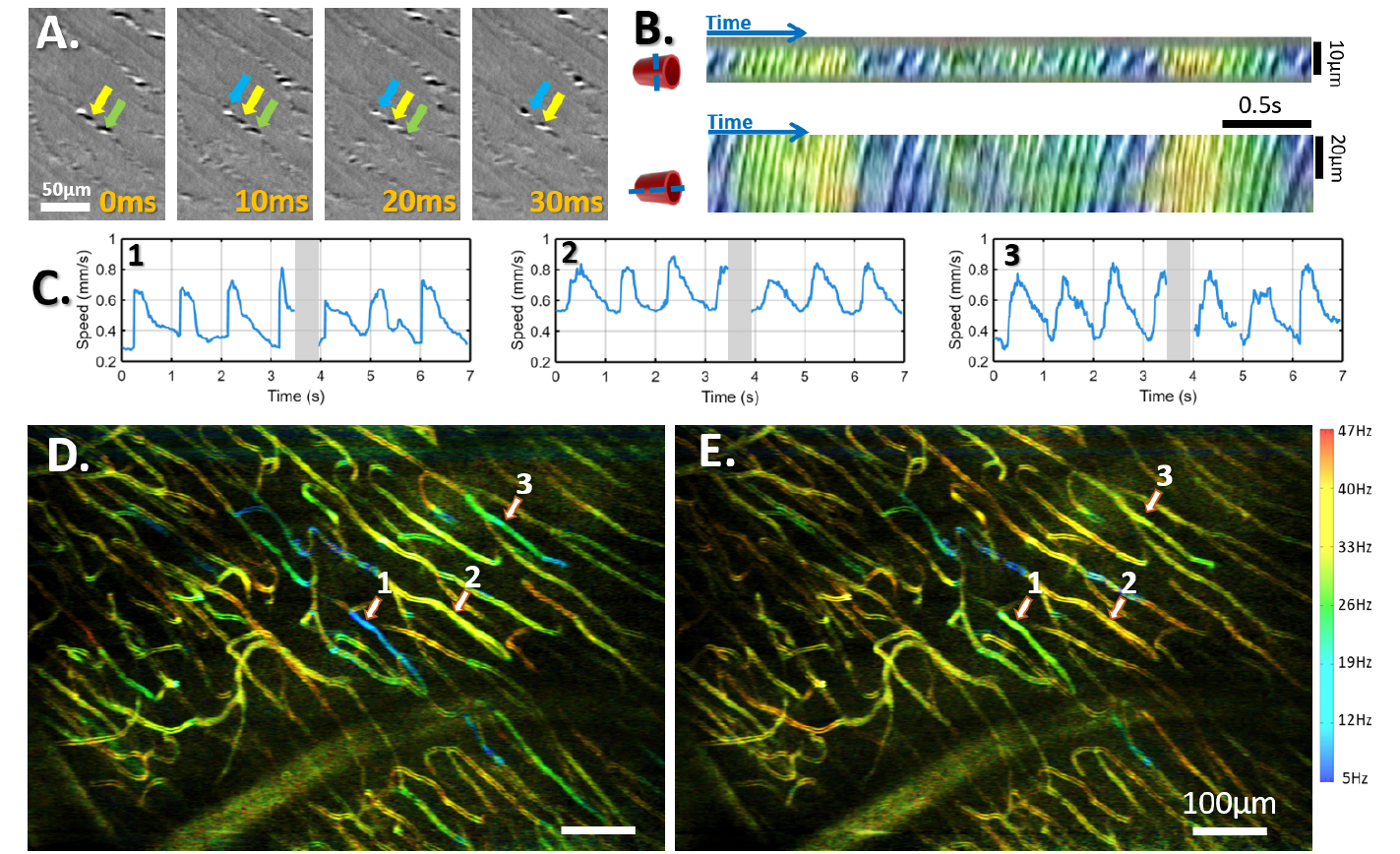
A. Time lapse representing four consecutive cropped phase-contrast images, where one can follow 3 individual red blood cells outlined respectively by blue, yellow, and green arrows. B. On the top, profile over time of a line perpendicular to a capillary, where individual red blood cells which went through the capillary can be tracked. Based on this information, we can compute the red blood cells rate (or frequency), which is color coded. B. on the bottom, profile over time of a line parallel to a capillary, where kymographs can be generated and blood flow speed can be computed. C. Blood flow speed over 7s (only interrupted by a blink represented by gray area) of three different capillaries indicated in D. D. and E. Color-coded perfusion map, where color represents the red blood cell frequency. Perfusion maps in D. and E. were generated with images acquired when the blood flow was minimal and maximum respectively. One can see that all capillaries change color, from blue/green to a more yellow/red aspect.

When assessing the temporal profile along the capillary, kymograph images can be generated (Fig. 7-B, bottom), where the angle between the axes provides the RBC speed. In this case, the speeds range from 0.3 mm/s to 0.8 mm/s, corresponding to the cardiac cycle, which is consistent with the range of speeds reported by previous studies [26, 27, 52]. The correlation between the frequency (color coded) and the angular component of the kymograph is evident. Figure 7-C illustrates the variation in blood flow speed over time for three distinct capillaries, which can be identified in Figures 7-D and E. Notably, the pulsatile nature of blood flow, reflecting the systole and diastole phases of the cardiac cycle, is clearly visible.

These analyses can be extended across the large FOV of AO-RSO images, as shown in Figs. 7-D and E and **Supplementary Figs. 1 and 2** for two additional subjects. Perfusion maps, color-coded according to RBC frequency, were generated using data collected over several cardiac cycles, specifically at time points corresponding to minimum and maximum speeds. **Visualizations 2 and 3** further showcase color-coded phase-contrast and motion-contrast videos, illustrating the AO-RSO system’s capacity to quantify blood flow both locally (at the level of individual capillaries) and across the entire FOV at a high frame rate. Notably, color variations synchronized with the cardiac cycle can be observed.

## 4. Discussion

We introduced the AO-RSO imaging system, a novel approach that uniquely combines the strengths of full-field and point-scanning detection. When imaging the retina in a clinical environment, gold-standard imaging modalities such as OCT, SLO, and fundus camera rely on very large FOV imaging to allow a global information to be evaluated for a faster identification of abnormalities. However, these modalities lack the cellular resolution needed for detailed local assessment that has proven to be crucial for early disease detection and precise therapy follow-up when using narrow FOV AO-SLO [50, 51, 53]. The AO-RSO method represents a crucial step towards bridging this gap by integrating wide FOV, cellular resolution, multimodality, and high frame rates within a single system, allowing for simultaneous acquisition of both global and localized retinal information. By reducing the need for multiple acquisitions to cover the required FOV in the clinical environment, the AO-RSO also improves patient comfort and streamlines clinical processes, facilitating clinical tasks such as generating cone density maps (Fig. 3-B), creating ultra-wide FOV montages (Fig. 6-B,C), and producing color-coded through-focus images to explore the 3D vascular network organization (Fig. 6-A). Furthermore, the high frame rate capability of the AO-RSO not only improves the efficiency of structural imaging, but also unlocks the potential to investigate neurovascular dynamics in the living human retina [5]. These dynamics are essential for understanding the interplay between nervous and vascular systems, with applications in identifying biomarkers for retinal, systemic, and neurodegenerative diseases. We further demonstrated the AO-RSO’s capability for multimodal imaging, integrating near-confocal bright-field detection with off-axis (or oblique illumination) detection for phase contrast imaging. These complementary modalities reveal diverse retinal structures: bright-field imaging highlights highly scattering features such as cones, rods, and nerve fibers, while phase contrast imaging reveals translucent structures such as vessel walls, red blood cells, ganglion cells, and photoreceptor inner segments. Although many of these features have been previously visualized using multimodal AO-SLO or DMD-based AO-FIO, those methods suffered from limited FOV and/or low frame rates [15, 21, 31, 32]. The AO-RSO overcomes these limitations, producing distortion-free phase-contrast images at high frame rates, thereby enabling the reliable extraction of biomarkers related to cellular structure and dynamics over a large FOV. For example, our system successfully measured blood flow velocity across a wide FOV in a single image sequence (Fig. 7 and **Supplementary Figs. 1 and 2**), offering unprecedented potential for studying capillary interactions and blood flow propagation.

Unlike traditional AO-SLO systems, where off-axis detection parameters are fixed (with few exceptions using DMD [54]), the AO-RSO system allows dynamic adjustment of the width and offset of the slit during imaging. This flexibility allows for optimization of the phase contrast for specific retinal features, accommodating variations in imaging requirements, such as for photoreceptor inner segments, red blood cells, and ganglion cells (see Fig. 5). The ability to adjust off-axis configurations is invaluable for imaging diverse retinal features and to advance our understanding of phase contrast mechanisms, including the effects of shape, size, refractive index, and feature depth. In this context, while imaging the RBCs, we observed additional structures within the capillaries, distinguished by their differences in size, shape, contrast amplitude, or contrast inversion (opposite biphasic profile), as exemplified in **Supplementary Fig. 3**. These findings highlight the importance of further investigating phase-contrast mechanisms to enable the accurate labeling of cellular features based on phase-contrast information.

The principle of the AO-RSO modality — line illumination synchronized with rolling shutter detection — can be adapted to other imaging modalities, including interferometric devices such as Time-Domain Full-Field OCT, which is a very promising imaging modality for both in-vivo human retinal imaging [55–57] and microscopy [3]. This adaptation could enhance sensitivity by rejecting multiply scattered photons and enable multimodal imaging, combining complementary coherent ballistic photon detection with forward-scattered photon detection for phase-contrast imaging. Furthermore, the AO-RSO principle and benefits extends beyond retinal imaging; in microscopy, a large FOV combined with high-resolution imaging enables the study of larger cell populations, their interactions, dynamics, and metabolic processes, further broadening the scope of this innovative approach.

While AO-RSO offers many advantages, it does have some limitations. Image processing is more complex compared to AO-FIO, as strip-based registration algorithms are required to correct intra-frame motion [40]. However, the system’s high frame rate allows precise correction for micro-saccades, enabling recovery of nearly all acquired images except those affected by blink events, unlike AO-FIO where micro-saccade events often necessitate image discard. Unlike AO-SLO and DMD-based systems, AO-RSO does not currently enable the simultaneous acquisition of bright-field and phase-contrast images. These must be collected sequentially, which reduces the frame rate for each modality, though still offering higher throughput than these systems. This limitation could be mitigated by integrating additional cameras dedicated to specific imaging modes or adopting faster cameras.

Another limitation lies in the directional nature of phase-contrast imaging, which, in the case of AO-RSO, is restricted to the vertical axis. Studies have shown that the off-axis detection orientation significantly influences phase contrast [18, 54, 58], with certain vessel walls disappearing when aligned perpendicular to the detection direction. In the AO-RSO system, while the walls of vertically oriented vessels are not visible in phase-contrast images, they can still be observed in perfusion maps, as indicated by the yellow arrows in Figs. 6 B and C.

Despite these constraints, our approach achieves a notable advancement by extending the capability to perform high-contrast, multimodal, cellular-resolution imaging of the human retina in-vivo over a wide FOV and at a high frame rate. This unique combination within a single system represents a pivotal advancement in retinal imaging, offering unprecedented capabilities for observing central nervous system dynamics in the living human body. Applications such as functional neuronal imaging [4] and the characterization of neurovascular coupling [5] are particularly promising, offering potential biomarkers for the early detection of retinal and neurodegenerative diseases, including Alzheimer’s disease [11].

## Conclusion

We have introduced the AO-RSO imaging modality, a novel system that combines the strengths of full-field and point-scanning techniques to deliver high-resolution, high-contrast, distortion-free images with a wide field of view and high frame rates. By utilizing line illumination synchronized with the rolling shutter detection of a 2D camera, AO-RSO enables precise spatial selection of detected photons: back-scattered photons for near-confocal bright-field imaging and forward-scattered photons for phase contrast imaging of translucent retinal features.

This modality has been successfully applied to in-vivo human retinal imaging, allowing visualization of diverse retinal structures, including photoreceptors, rods, nerve fiber bundles, red blood cells, vessel walls, and ganglion cells, across a wide FOV and spanning various retinal eccentricities, from the fovea to the periphery. Furthermore, AO-RSO’s ability to dynamically adjust off-axis detection enhances phase contrast optimization for specific retinal features, highlighting its versatility and adaptability.

We have also demonstrated the system’s capability to extract critical biomarkers over a large FOV, such as photoreceptor density and blood flow velocity, underscoring its potential for advanced clinical diagnostics and fundamental research. By addressing key limitations of existing modalities, AO-RSO provides a transformative approach to retinal imaging, opening new avenues in biomedicine for the study of structural details and cellular dynamics.

## 8. Supplemental document

### 8.1. Color-coded Perfusion Map from S3

**Fig. 8.**
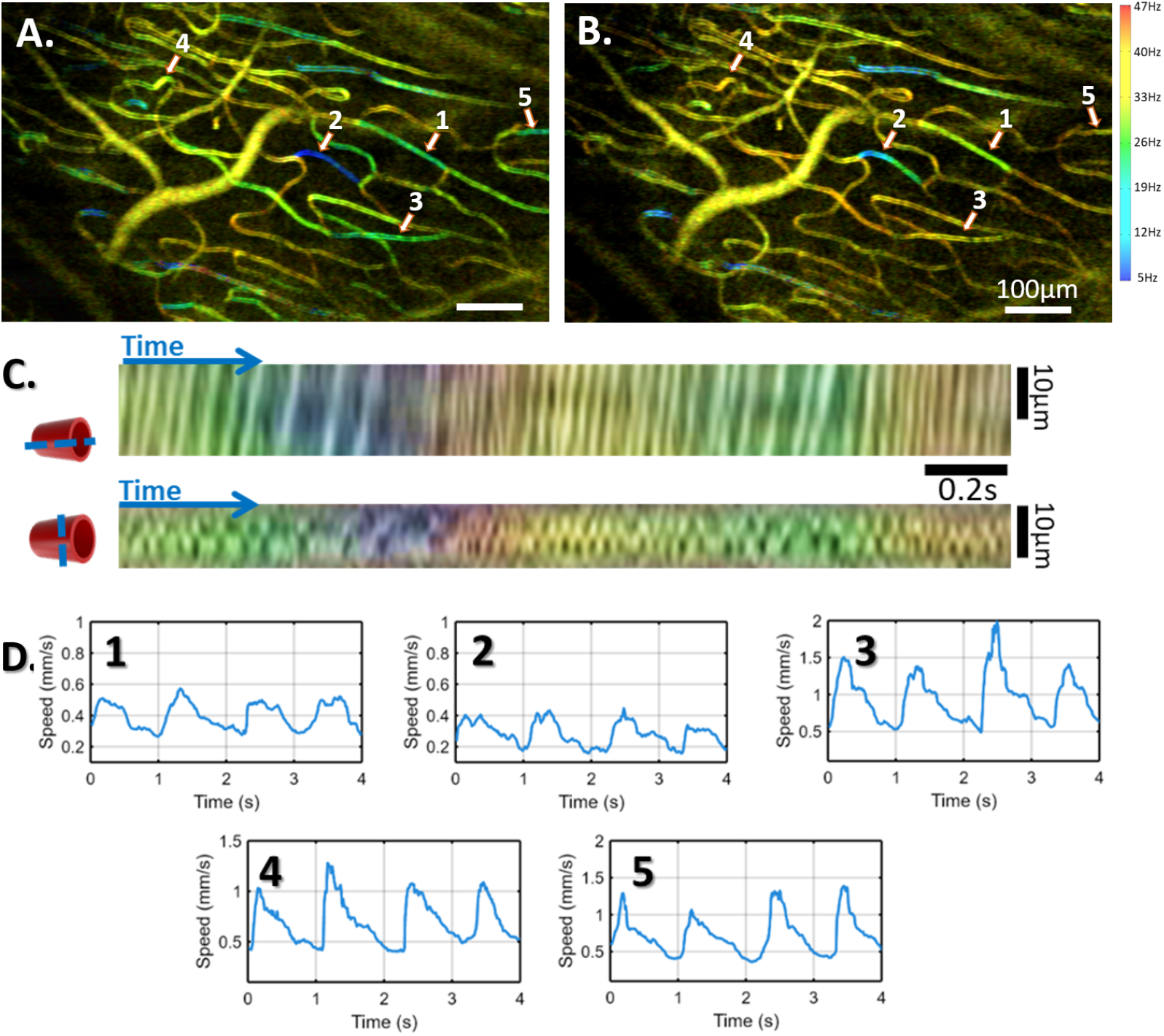
A. and B. Color-coded perfusion map, where color represents the red blood cell frequency. Perfusion maps in A. and B. were generated with images acquired when the blood flow was minimal and maximum respectively. C. Red blood cells tracing over time within (top) and perpendicular (bottom) to the capillary. D. Blood flow speed over time for capillaries 1 to 5 indicated in A. and B.

**Fig. 9.**
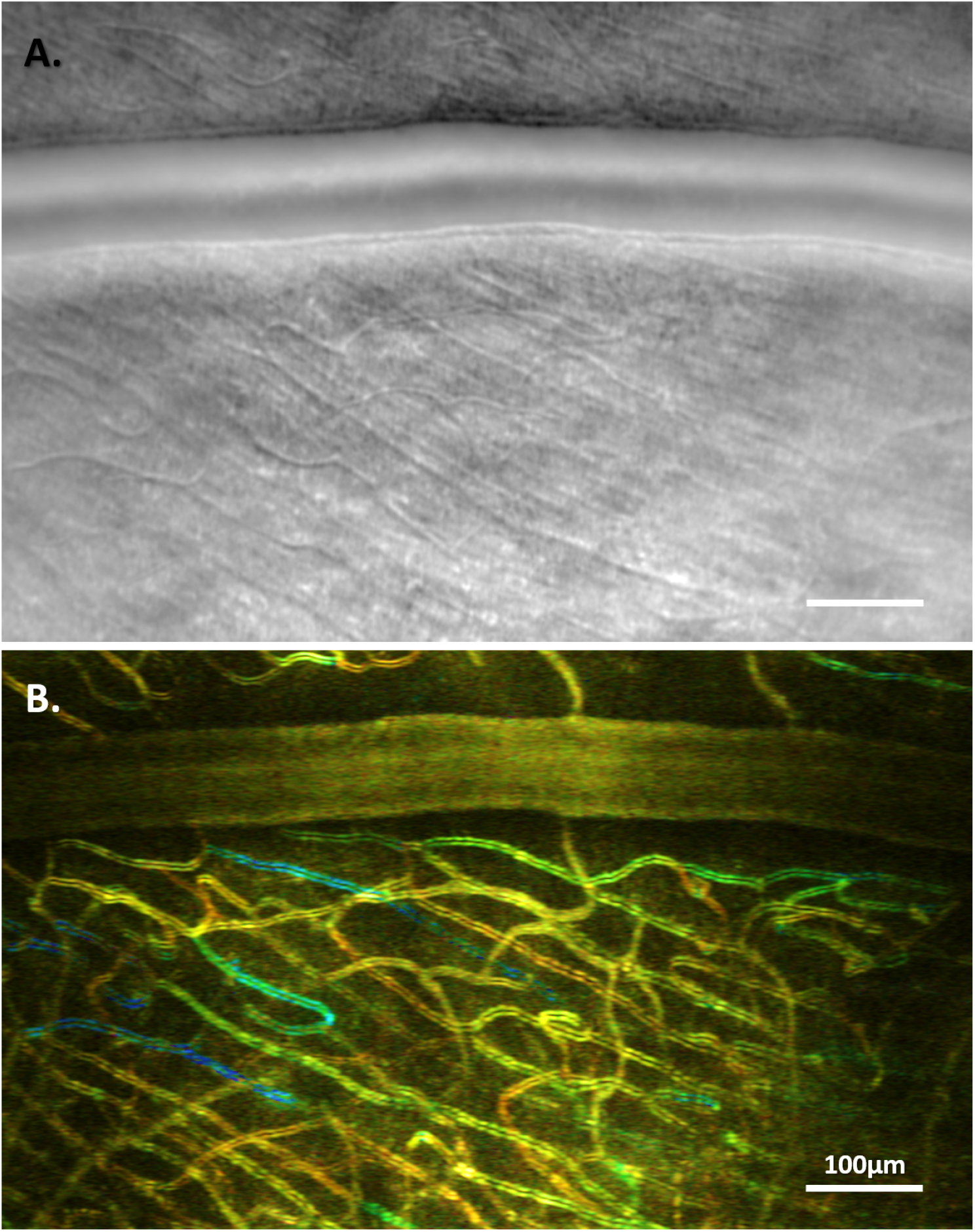
A. Cropped phase-contrast image. B. Color-Coded Motion Contrast Perfusion Map for an averaged frequency over time.

**Fig. 10.**
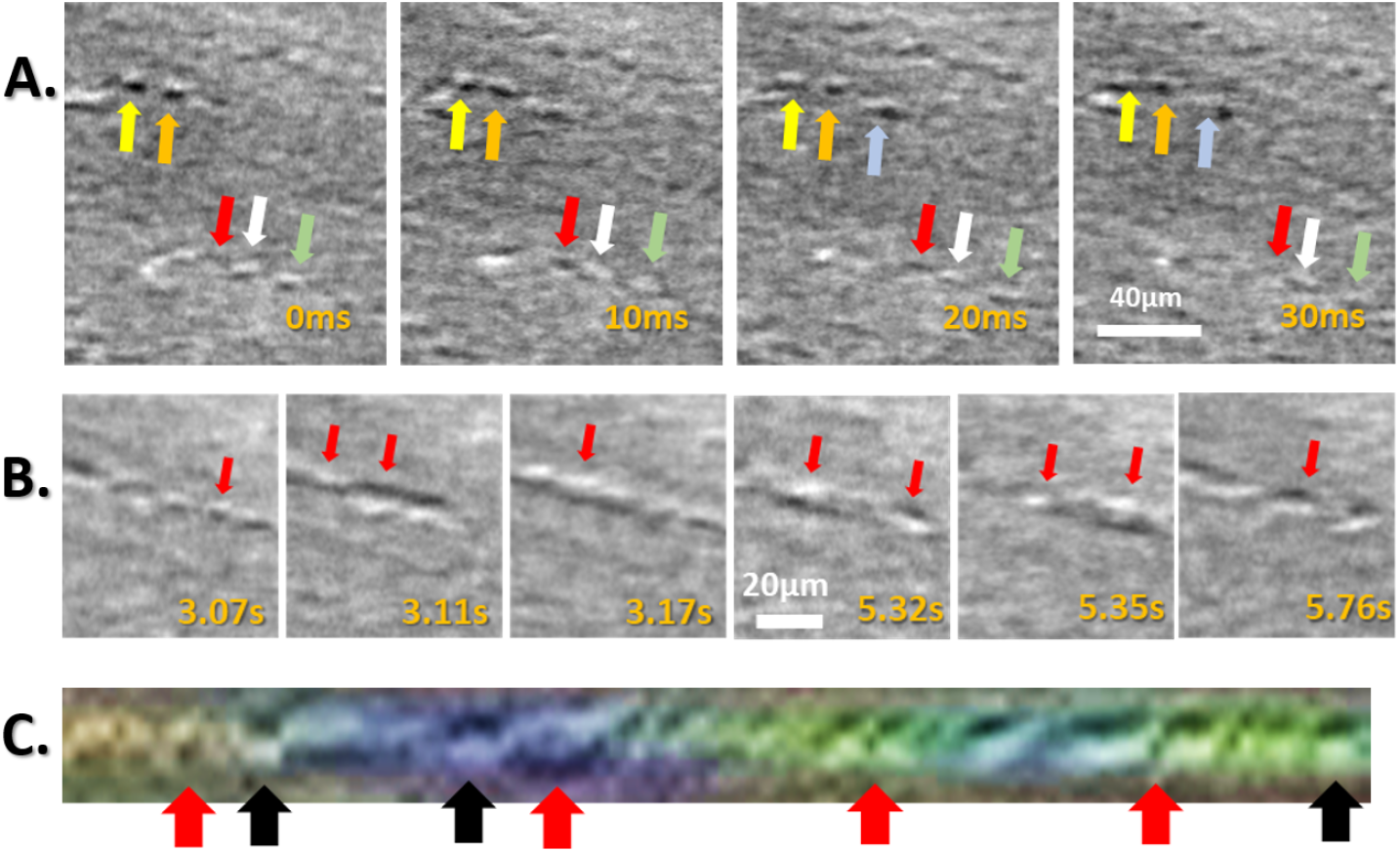
A. Cropped phase-contrast consecutive images, where one can follow individual red blood cells at different time points and outlined by arrows of the same color. Note that the red arrow outlines a cellular structure within the capillary with inverted contrast compared to the others. B. Cropped phase-contrast images of the same capillary for different time points (indicated on the bottom right). Red arrows indicate different structures flowing inside the same capillary with different size, shape, contrast amplitude and inverted contrast. C. tracing over time of a perpendicular line over the capillary in B. showing different cellular structures in terms of size, shape, speed, contrast amplitude and contrast polarization flowing inside the capillary. Note that features indicated by red arrows present an inverted contrast compared to those highlighted by the black arrows.

### 8.2. Color-coded Perfusion Map from S4

### 8.3. Contrast inversion and different cellular features within capillaries

### 8.4. Visualization 1

Visualization 1: Photoreceptor imaging close to the foveal center at high frame rate (200Hz) and over a wide FOV (4.5*°* × 2.5*°*). Individual cone photoreceptors can be tracked over time throughout the extension of the wide FOV without any motion-induced image distortion.

### 8.5. Visualization 2

Visualization 2: Phase contrast video acquired at 100Hz and displayed here at 25Hz, where color codes for red blood cells frequency. Note that the AO-RSO can reliably quantify blood flow over a wide FOV and for each visible individual capillary.

### 8.6. Visualization 3

Visualization 3: Same sequence as Visualization 2 but here showing the Perfusion Maps computed for 20 consecutive images where the color codes for red blood cell frequency.

